# STonKGs: A Sophisticated Transformer Trained on Biomedical Text and Knowledge Graphs

**DOI:** 10.1101/2021.08.17.456616

**Authors:** Helena Balabin, Charles Tapley Hoyt, Colin Birkenbihl, Benjamin M Gyori, John Bachman, Alpha Tom Kodamullil, Paul G Plöger, Martin Hofmann-Apitius, Daniel Domingo-Fernández

**Author notes:** **Corresponding Author:** Balabin, H., and Domingo-Fernández, D. Department of Bioinformatics, Fraunhofer Institute for Algorithms and Scientific Computing (SCAI), Sankt Augustin 53757, Germany. Telephone details. and.

## Abstract

The majority of biomedical knowledge is stored in structured databases or as unstructured text in scientific publications. This vast amount of information has led to numerous machine learning-based biological applications using either text through natural language processing (NLP) or structured data through knowledge graph embedding models (KGEMs). However, representations based on a single modality are inherently limited. To generate better representations of biological knowledge, we propose STonKGs, a Sophisticated Transformer trained on biomedical text and Knowledge Graphs. This multimodal Transformer uses combined input sequences of structured information from KGs and unstructured text data from biomedical literature to learn joint representations. First, we pre-trained STonKGs on a knowledge base assembled by the Integrated Network and Dynamical Reasoning Assembler (INDRA) consisting of millions of text-triple pairs extracted from biomedical literature by multiple NLP systems. Then, we benchmarked STonKGs against two baseline models trained on either one of the modalities (i.e., text or KG) across eight different classification tasks, each corresponding to a different biological application. Our results demonstrate that STonKGs outperforms both baselines, especially on the more challenging tasks with respect to the number of classes, improving upon the F1-score of the best baseline by up to 0.083. Additionally, our pre-trained model as well as the model architecture can be adapted to various other transfer learning applications. Finally, the source code and pre-trained STonKGs models are available at https://github.com/stonkgs/stonkgs and https://huggingface.co/stonkgs/stonkgs-150k.

## 1. Introduction

In recent years the availability of biomedical data has increased drastically (Dash *et al*., 2019). Such data originate from a vast collection of modalities such as high-throughput experiments, electronic health records as well as cell-based and biochemical assay data. The information derived from research carried out on those data is commonly stored in two distinct forms: 1) as unstructured free text in scientific publications, and 2) in condensed, structured biomedical networks. However, the biology represented in the literature strongly depends on the different contexts that it occurs in. For instance, certain proteins or chemicals may exclusively interact with others in a specific tissue or cell type (Stacey *et al*., 2018), or specific biochemical reactions may only take place under certain conditions. Consequently, in order to leverage the most out of the biomedical knowledge stored in both structured and unstructured formats, it is of utmost importance to study each relation in the relevant context it was observed in. While networks often lack this contextual information due to their inherent degree of abstraction (Saqi *et al*., 2019), unstructured text contains context at the expense of explicit logical structure. Thus, the complementary strengths from both sources could be leveraged to enable a more complete, context-specific, and actionable representation of biological knowledge.

Knowledge graphs (KGs) represent information in a structured manner in order to encode the broad spectrum of complex interactions occurring in biology. In order to exploit the information contained in KGs through machine learning algorithms, numerous knowledge graph embedding models (KGEMs) have been developed to encode the entities and relations of KGs in a higher dimensional vector space while attempting to retain their structural properties (Ji *et al*., 2021). Utilizing the resulting vector representations, more sophisticated tasks can be conducted (i.e., link prediction, node classification, and graph classification). When these KGs contain more detailed, contextualized descriptions of biological interactions, the performance of KGEMs can be substantially improved. Such improvements can be achieved by incorporating metadata that specifies the context of each relation (e.g., the pH value in which a molecular interaction occurs or the specific cell type in which a protein is expressed). Therefore, context-specific KGs have recently been used in combination with other data modalities in several biomedical applications. For instance, Federico and Monti (2021) demonstrated how to gain insights on specific human cell-line processes by annotating protein-protein interaction networks with contextualized cell-line information extracted from the scientific literature. Similarly, a recent study from Doncheva *et al*. (2021) introduced a methodology that proposes the most suitable organism to model a human pathway by evaluating whether the expression of genes in a certain pathway across four species (i.e., rat, mouse, pig and humans) is maintained in the same tissue. To achieve this, the authors leveraged a contextualized protein-protein interaction network generated with ortholog information together with transcriptomics data and mentions of proteins in the scientific publications.

Due to the availability and abundance of unstructured text data in scientific literature and electronic health records, natural language processing (NLP) has become an important tool for extracting information on biomedical contexts. Similar to KGEMs, language models (LMs) are used to transform their input, namely word sequences, into a high-dimensional vector space, resulting in so-called embeddings. One approach to learning these embeddings in a contextualized manner is through the use of the attention mechanism (Vaswani *et al*., 2017), which is for instance employed in the Bidirectional Encoder Representations from Transformers (BERT) model by Devlin *et al*. (2019). Its biomedical counterpart, BioBERT (Lee *et al*., 2020), is pre-trained on a large PubMed text corpus to learn a contextualized representation of biomedical knowledge. Such a pre-trained Transformer can then be used on a variety of classification tasks (e.g., named entity recognition (Sun *et al*., 2016), sequence classification (Baker *et al*., 2016) and question answering (Tsatsaronis *et al*., 2015)) with minimal model architecture adaptations in a so-called fine-tuning procedure. The goal is to leverage and flexibly adapt the pre-trained embedding representations, which is especially beneficial for fine-tuning tasks with small training datasets.

To incorporate other data modalities, Transformers with cross-modal attention have been proposed as an extension to purely text-based Transformer models. For instance, Tsai *et al*. (2019) used cross-modal attention to capture complex interdependencies between text, video, and audio data to enhance the frame of reference of context-specific LMs. More recently, Kamath *et al*. (2021) improved state-of-the-art performances on multiple visual reasoning tasks by applying a cross encoder on a concatenation of textual and visual embeddings. Moreover, several Transformer-based LMs have demonstrated the benefit of incorporating structured KG data in the general (Zhang *et al*., 2019) as well as the biomedical domain (He *et al*., 2020; Fei *et al*., 2020). However, the former approaches operate at a word level (rather than sentence level) by combining textual embeddings from LMs and entity embeddings from KGs through entity linking (i.e., the process of aligning text tokens and KG entities). Recently, Sun *et al*. (2020) proposed a different strategy for combining information from KGs and text by concatenating word, entity, and relation embeddings at the sentence level. Similarly, Nadkarni *et al*. (2021) have combined textual descriptions of nodes with embedding representations learned by KGEMs for link prediction. Finally, Transformer-based LMs have also been directly applied on graph-structured data (Ying *et al*., 2021).

Here, we present STonKGs, a Sophisticated Transformer trained on biomedical text and Knowledge Graphs. STonKGs is a multimodal approach that combines subgraph-level information from a KG with corresponding sentence-level text data from literature to learn better embedding representations. We demonstrate STonKGs on a KG consisting of millions of text-triple pairs extracted from the biomedical literature and pathway databases, assembled using the Integrated Network and Dynamical Reasoning Assembler (INDRA) (Gyori *et al*., 2017). Using this dataset, we benchmark STonKGs against two baseline models (i.e., BioBERT (Lee *et al*., 2020) and node2vec (Grover and Leskovec, 2016)) in a transfer learning setting on eight different fine-tuning tasks corresponding to distinct biological applications. Our results highlight how combining both modalities can enable STonKGs to outperform both baselines, particularly on more challenging classification tasks with a larger number of classes. Furthermore, the STonKGs model architecture can be easily adapted to other applications on text-triple pairs in the biomedical as well as general domain. We released the source code and pre-trained STonKGs models at https://github.com/stonkgs/stonkgs and https://huggingface.co/stonkgs/stonkgs-150k.

## 2. Methods

Our main goal was to evaluate the effect of combining text and KG data in the proposed model architecture (i.e., STonKGs). As a data resource, we used the INDRA KG, which contains millions of triples with text evidence and annotations, further described in Section 2.1 (**Figure 1A**). We compared our proposed STonKGs model against two baseline models which only used one of the respective knowledge sources in a unified experimental setting (**see Section 2.2 and Figure 1B**). Next, we outline our evaluation setting consisting of eight different classification tasks (**Section 2.3**). Finally, we describe the software implementation and hardware used to conduct this work in Section 2.4.

**Figure 1.**
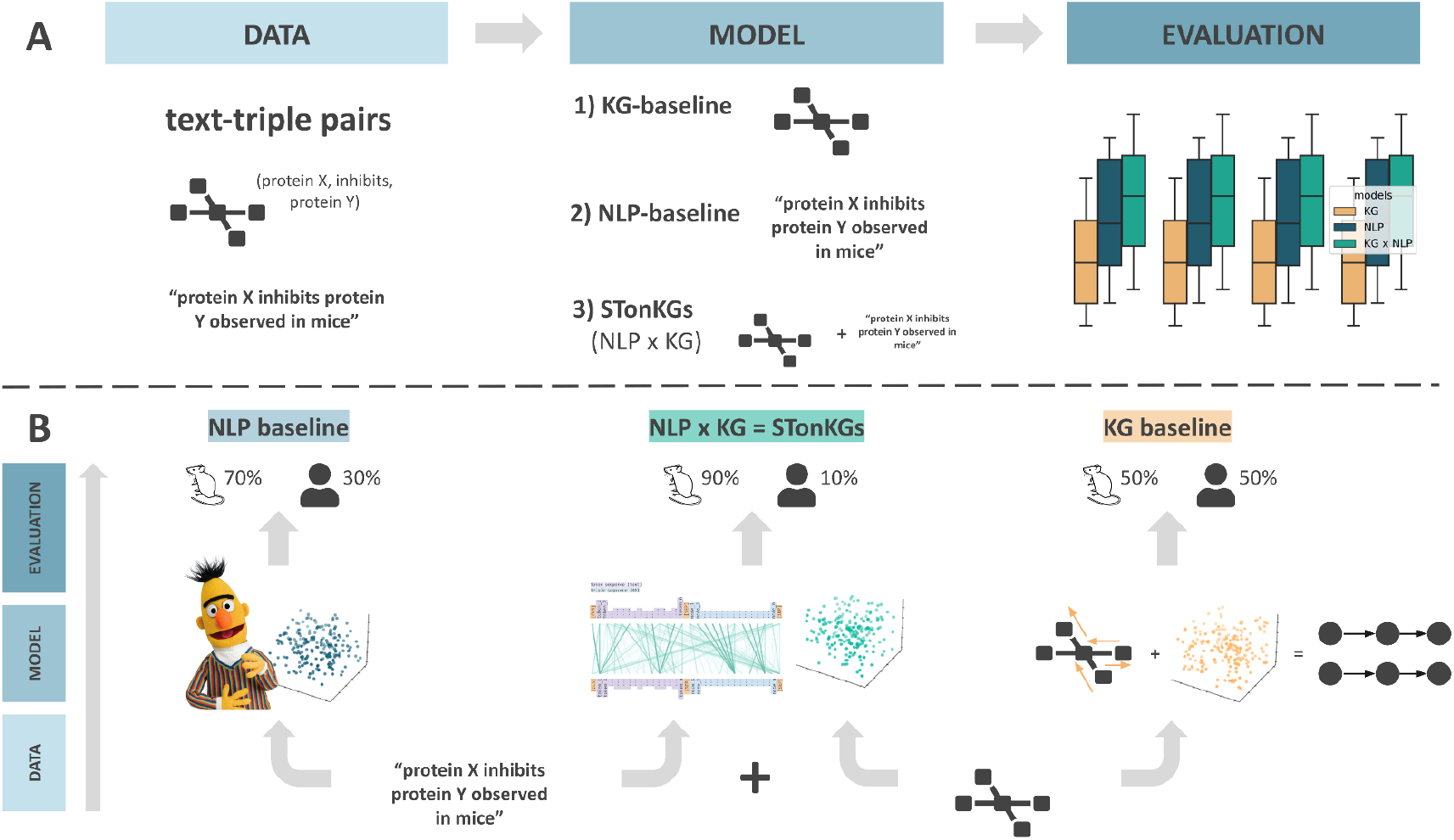
Methodology workflow. This figure illustrates the classification of the context annotation for a given text-triple pair. In this example, the models aim to predict the species in which a certain biological process was observed (e.g., mice). **A)** The three models (i.e., the two baselines and the proposed STonKGs model) are trained and evaluated in a shared experimental setting. **B)** For each text evidence and triple pair, the two baseline models exclusively employ a single modality, whereas STonKGs leverages both.

### 2.1. Dataset

To combine the structured information represented in a KG with unstructured text, we required a KG containing relations for each triple and the corresponding text evidence from which the triple has been extracted. As a result, our dataset consisted of text-triple pairs such as (“*Sorafenib is a multi-kinase inhibitor that inhibits various kinases including VEGFR-2*”, **(Sorafenib, directlyDecreases, VEGFR-2**), which is represented as **(a(pubchem.compound:216239), directlyDecreases, kin(p(hgnc:6307))**). We employed a KG containing 35,150,093 triples assembled by INDRA (Gyori *et al*., 2017) from pathway databases and the output of text mining systems **(Supplementary Table 1)** run on i) MEDLINE abstracts, ii) PubMed Central full text articles, and iii) several publishers’ text mining corpora **(see Supplementary Figure 1 for details on node and relation types)**. The original version of the INDRA KG comprised non-grounded nodes (i.e., nodes that could not be normalized to a standardized ontology) and triples without text evidence, both of which were filtered out in a preliminary data cleaning step **(described in Supplementary Text 1)**. Ultimately, the preprocessed version of the INDRA KG consisted of 174,534 nodes and 13,609,994 triples. Out of all triples, 127,149 were selected for each of the eight fine-tuning tasks since they have been labelled with context-specific information (i.e., annotation class) or they have been manually curated for classification tasks **(see Supplementary Table 2)**. The 13,482,845 remaining non-annotated triples were used in the unlabelled pre-training procedure **(see Supplementary Text 1)**.

### 2.2. Models

As shown in **Figure 1**, all three models were operating under the same experimental conditions (i.e., in the same transfer learning setting, evaluated on the same tasks), with the exception of their utilized modalities. In contrast to the NLP- and KG-baselines (i.e., text evidence and triple-based models), STonKGs jointly builds upon both modalities. The following subsections outline our proposed STonKGs model architecture as well as the two baseline models used as a benchmark. All three models shared the same two-fold training procedure, consisting of a pre-training and a fine-tuning part. The architectural change in the fine-tuning procedure was equivalent across all three models and consisted of placing a classification head on top of the pre-trained model.

#### 2.2.1. NLP-baseline

The NLP-baseline was built on the pre-trained BioBERT v.1.1 model (Lee *et al*., 2020), a Transformer-based language model trained for 1 million steps on chunks of 512 tokens from a 4.5 billion token corpus stemming from PubMed abstracts **(see Supplementary Table 3 for an overview on the hyperparameters of this model)**. To prepare text evidences from INDRA statements for the NLP-baseline, the contiguous string of text was first split into single (sub)words (i.e., tokens), using the pre-trained tokenizer of BioBERT. The resulting token sequence was extended with special classification and separator tokens (i.e., [CLS] and [SEP]), and then padded or truncated accordingly to match the fixed input length of the language model (512 tokens, i.e., a paragraph). Passing the sequence through BioBERT yielded token embedding vectors for a given text evidence, in which each of these embedding vectors is based on the weighted average of its surrounding tokens that is learnt by the attention mechanism of a pre-trained Transformer (Vaswani *et al*., 2017). This procedure ensures that each token embedding vector contains the context of its surrounding tokens.

In order to adapt BioBERT as a classifier for text evidences in a fine-tuning procedure, further model components, namely, pooling and a final linear layer with a softmax activation function, were added to enable sequence classification. In line with a commonly used aggregation technique derived from Devlin *et al*. (2019), our pooling procedure consists of using the special classifier (i.e., [CLS]) token embedding vector as a representation of the overall token embedding sequence for a given text evidence. This token embedding vector is used as an input for the final linear layer to generate class probabilities for the provided text evidence. Finally, we would like to note that in this transfer learning setting, we not only trained the parameters of the sequence classification components, but also fine-tuned all parameters of the entire model architecture, including the weights of the BioBERT model.

#### 2.2.2. KG-baseline

The input for the KG-baseline are high-dimensional node embeddings learnt by node2vec (Grover and Leskovec, 2016) using the hyperparameters listed in **Supplementary Table 3**. Similar to the embeddings of word sequences produced by word2vec (Mikolov *et al*., 2013), node2vec generates embeddings for node sequences based on random walks. As a result, the embedding of a given node is formed based on the structure of its surrounding network neighborhood.

In concordance with the other two models, our KG-baseline relied on sequential inputs for each triple. Therefore, we designed a novel approach that generated a sequential representation for each triple while incorporating the embeddings generated by node2vec **(see Figure 2)**. The general idea behind the sequential representation is to generate a sequence of embeddings *e*(*h*_*i*_,*t*_*i*_) for the two nodes *h*_*i*_,*t*_*i*_ in the i-th triple (*h*_*i*_,*r*_*i*_,*t*_*i*_). To do so, our proposed approach leveraged the sequence of random walks *h* = (*h*_*i*_, …,*h*_*n*_) and *t* = (*t*_*i*_, …,*t*_*n*_) generated by node2vec for *h*_*i*_ and *t*_*i*_, replacing each random walk by the embeddings 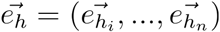 and 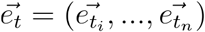 learnt for each node in the walk. Subsequently, we acquired the embedding sequence of a given triple as the concatenation of the random walk-based embedding sequences of its two nodes 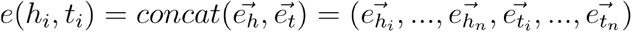. This final random walk-based sequential representation, as opposed to other alternatives (e.g., concatenation of the two original node embeddings of a given triple), ensured a fair comparison, since the other two models (i.e., NLP baseline and STonKGs) are also based on sequential inputs.

**Figure 2.**
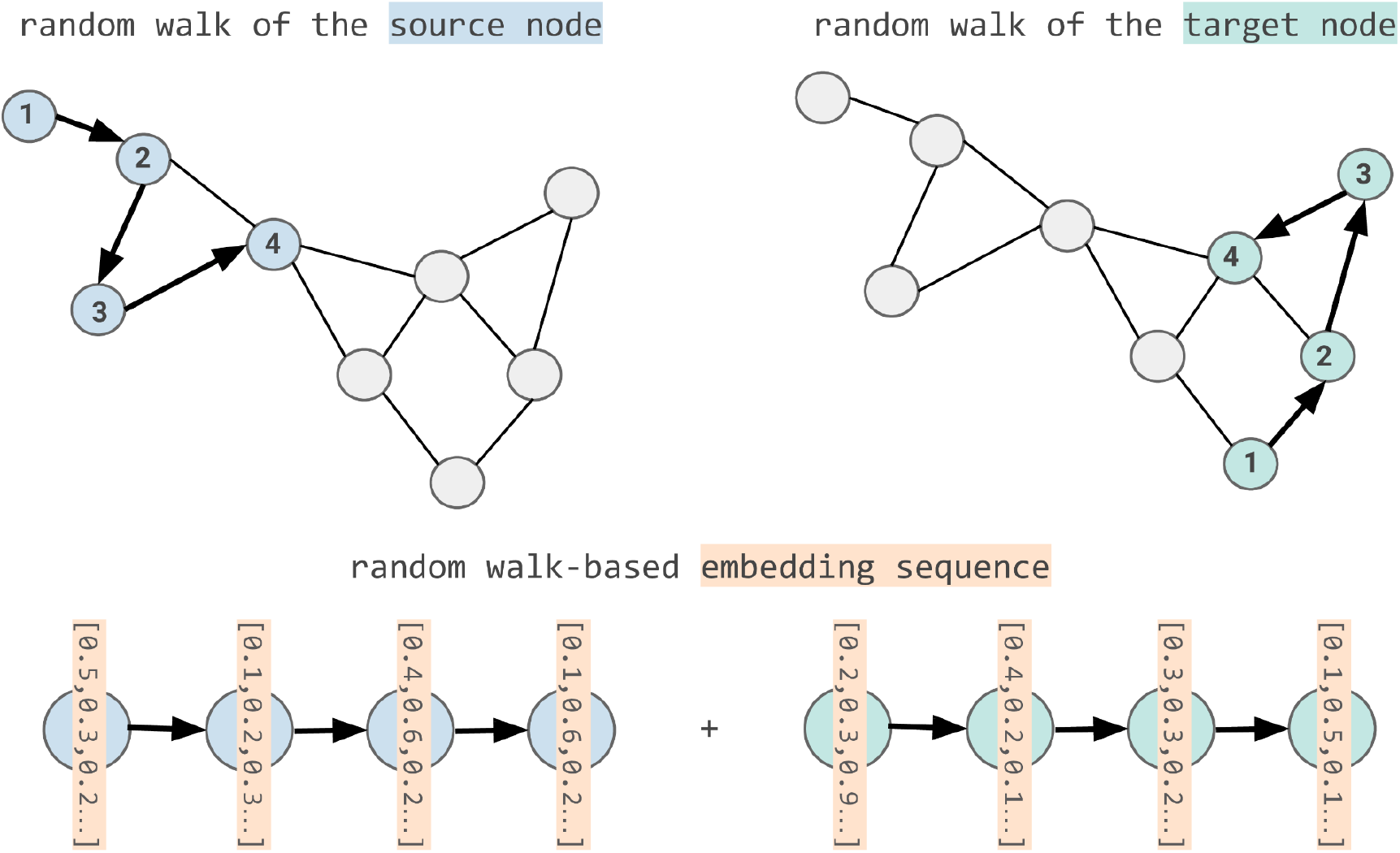
Transforming KG embeddings into sequential inputs. For a given triple (*h*_*i*_,*r*_*i*_,*t*_*i*_), we generate the final random walk based embedding representation *e*(*h*_*i*_,*t*_*i*_) based on the following steps: 1. Obtain the random walks based on the pre-trained node2vec model: *h* = (*h*_*i*_, …,*h*_*n*_) and *t* = (*t*_*i*_, …,*t*_*n*_) for *h*_*i*_ and *t*_*i*_ 2. Embed each node in those random walks, resulting in two random walk-based embedding sequences: 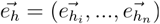 and 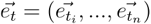 3. Generate the final embedding sequence 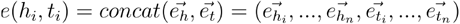

Similar to the NLP baseline outlined in Section 2.2.1, the embedding sequences for each triple are pooled, and passed through a linear layer with a softmax activation function to generate the final classification labels. Here, the pooling operation is defined as the dimension-wise maximum of the sequence embeddings, consequently mapping the sequence to a single vector. Since the KG-baseline employs static embeddings for the final classification task, the KG-baseline did not technically fit into the pre-training and fine-tuning paradigm used in NLP. However, for the sake of consistency, we will refer to the feature extraction based on transfer learning (i.e., embeddings from node2vec) used in the KG-baseline as pre-training and the final classification tasks as fine-tuning procedures as well.

#### 2.2.3. STonKGs

Similar to BERT, STonKGs consists of multiple stacked Transformer layers with the attention mechanism forming the core of the overall model architecture. However, in contrast to the standard attention mechanism applied on text tokens, STonKGs uses a joint Transformer on a concatenation of text tokens and KG nodes, as illustrated in **Figure 3**. In accordance with the terminology introduced by Kamath *et al*. (2021) for their joint Transformer (on image and text data), this Transformer is hereafter referred to as a cross encoder. The rationale behind using a cross encoder over other information fusion techniques was that it allows for learning implicit alignments between text tokens and KG nodes without requiring any entity linking step between the two modalities. More specifically, the interdependencies in the combined input sequence are represented by attention weights, shown by the links between the inputs in **Figure 3**. These weights are learnable parameters used to calculate weighted average representation of a given entity (i.e., text token or node of the input sequence) based on the embedding vectors of its surrounding entities from both modalities. As a result, the calculated representation of each entity contains contextual information from the KG and text input.

**Figure 3.**
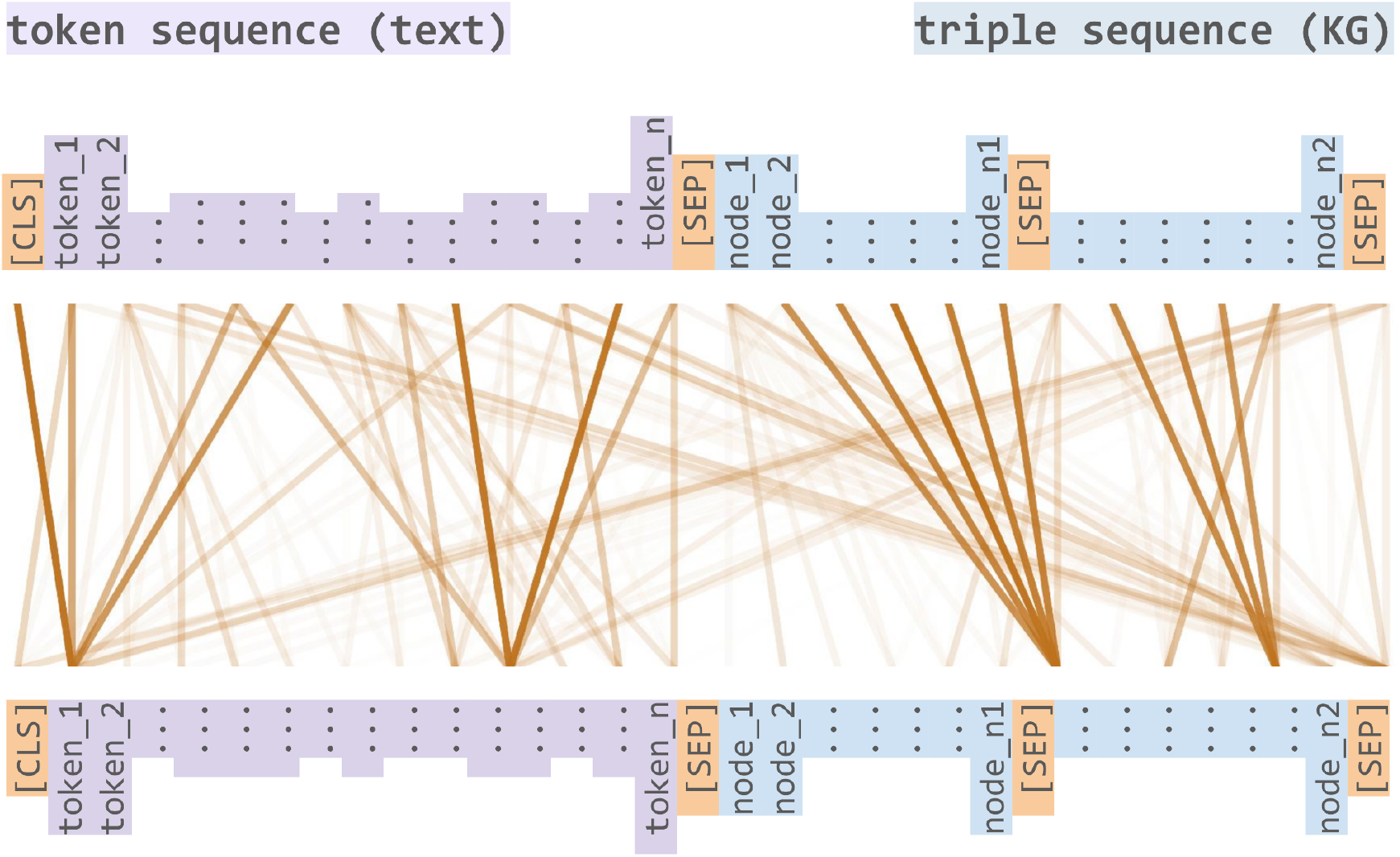
Cross-modal attention between text data (token sequences) and KG data (triple sequences). The input is a concatenation of a token and a triple sequence. Each element in the initial input sequence consists of its respective BioBERT embedding. The resulting hidden states are processed by two different heads for text tokens and KG nodes, respectively. While the MLM head is returning probabilities for each token of the NLP-backbone, the MEM head is converting the hidden states onto probabilities for each node of the KG-backbone.

To construct the cross encoder of STonKGs, we used the same hyperparameters as the BERT_BASE_ model (see Devlin *et al*., 2019), such as the maximum sequence length (512 tokens), hidden state dimension, number of Transformer layers, and attention heads. We used embeddings of the combined text and KG input sequences as the input to STonKGs, based on the text-triple pairs extracted from the INDRA statements. The overall input sequence length was split into half, in order to comprise 256 text tokens and 256 KG nodes (including special tokens). The initial embedding sequences of the text-triple pairs were generated with BioBERT and node2vec for text and triples, respectively, which we will refer to as the NLP- and KG-backbone in the following (based on the steps outlined in Section 2.2.1 and 2.2.2). However, instead of simply concatenating the random walk-based embedding sequences of the two nodes of a triple, we further added a [SEP] token between and after the two random walk sequences 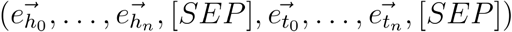, as shown in **Figure 3**. The use of the special separator token intends to structurally differentiate between text and KG data in the input sequence, similar to the distinction of two input sentences in the original BERT model. Moreover, we masked some of the input using the embedding vector of the special [MASK] token from the NLP-backbone (the masking strategy is explained in detail below). Additionally, we used positional and segment embeddings to further distinguish text and KG nodes of the combined input sequence in our cross encoder. Given the described inputs of STonKGs, the model has three different training objectives during pre-training, which are jointly used to learn the parameters of the cross encoder:

1. **Masked Language Modeling (MLM)**: For the first 256 text tokens, we employed the same MLM task and followed the same masking procedure used in the pre-training process of BERT. The goal of this task is to correctly predict the masked tokens based on a so-called MLM head. This head consists of a linear layer followed by the softmax function, which maps the final hidden states of the cross encoder to probabilities for each token in the vocabulary of the NLP-backbone.
2. **Masked Entity Modeling (MEM):** Inspired by the original MLM task, we built a counterpart for predicting masked nodes for the latter half of the combined input sequence (i.e., the KG input), again using the same masking strategy as in BERT. In this case, the goal is to correctly predict masked nodes in the random walk-based embedding sequences. Analogous to the MLM head, our custom MEM head consists of a linear layer followed by a softmax function. However, unlike the MLM head, the MEM head maps the hidden states to probabilities for each node occurring in the KG of the KG-backbone (as well as the [SEP] token, to remain consistent with the added [SEP] tokens) **(see Figure 3)**.
3. **Next “Sentence” Prediction (NSP):** Similar to the original NSP task, we designed an equivalent training objective that aims to correctly predict whether a text and triple belong to each other, or whether they are randomly chosen from distinct INDRA statements. In accordance with Devlin *et al*. (2019), we also used the final hidden state of the [CLS] token for this binary prediction task. However, in order to preserve as much of the original training data as possible, we decided to augment the training data (rather than replace entries in it) with negative samples. In our case, we used 25% of the original pre-training dataset size, which is significantly smaller than the 50% used in BERT.

As a result, the pre-training objective of STonKGs consists of minimizing the total loss, more specifically, the sum of the losses across all three training objectives: ℒ_total =_ ℒ_MLM_ + ℒ_MEM_ + ℒ_NSP_. All relevant hyperparameters used for the pre-training process of STonKGs (e.g., batch size and learning rate) are listed in **Supplementary Table 3**.

In order to evaluate STonKGs on each of the eight fine-tuning tasks (explained in the next section), we followed the same procedure that is outlined in Section 2.2.1 (NLP-baseline). Consequently, we employed a classification head on top of the pre-trained STonKGs architecture, consisting of a pooling step, a linear layer, and a softmax activation function to generate class probabilities for a given text-triple pair. Similar to the NLP-baseline, we also utilized the [CLS] token for pooling, and tuned all parameters of the entire STonKGs model architecture in our fine-tuning tasks.

### 2.3. Evaluation

In line with other Transformer-based transfer learning approaches, we used the majority of the INDRA text-triple pairs, predominantly unannotated triples, for pre-training **(see Section 2.1**), and the remaining annotated text-triple pairs (approximately 1.63%) were used for the fine-tuning datasets. We evaluated the models on a benchmark consisting of eight fine-tuning tasks, namely, two relation type classification tasks, four context annotation tasks, and two correct/incorrect tasks (tasks 1-2, 3-6, 7-8 in **Table 1**, respectively). The relation type tasks consist of two binary classifications in which each model either predicts the polarity (i.e., increase or decrease) or the type of interaction (i.e., direct or indirect interaction) of a given triple. The four context annotation tasks aim to predict the class (i.e., the context) of given text-triple pairs in a variety of biomedical settings: i) cell line, ii) disease, iii) cellular location, and iv) species. All of these cases represent multiclass classification tasks employing between three and ten classes depending on the most common occurrences of classes in each of the contexts. Finally, the two correct/incorrect tasks consist of a binary classification task where the model determines whether the text-triple pair is correct or incorrect, and a multiclass task where the model not only determines whether it is correct or incorrect but also which type of error it is. The sample sizes of the task-specific fine-tuning datasets ranged from 3,760 to 78,979 text-triple pairs, depending on the availability of triple annotations. An overview on the tasks as well as their respective summary statistics can be found in **Supplementary Table 2**. The distribution of classes of the fine-tuning tasks can be found in **Supplementary Figure 2**.

**Table 1.**
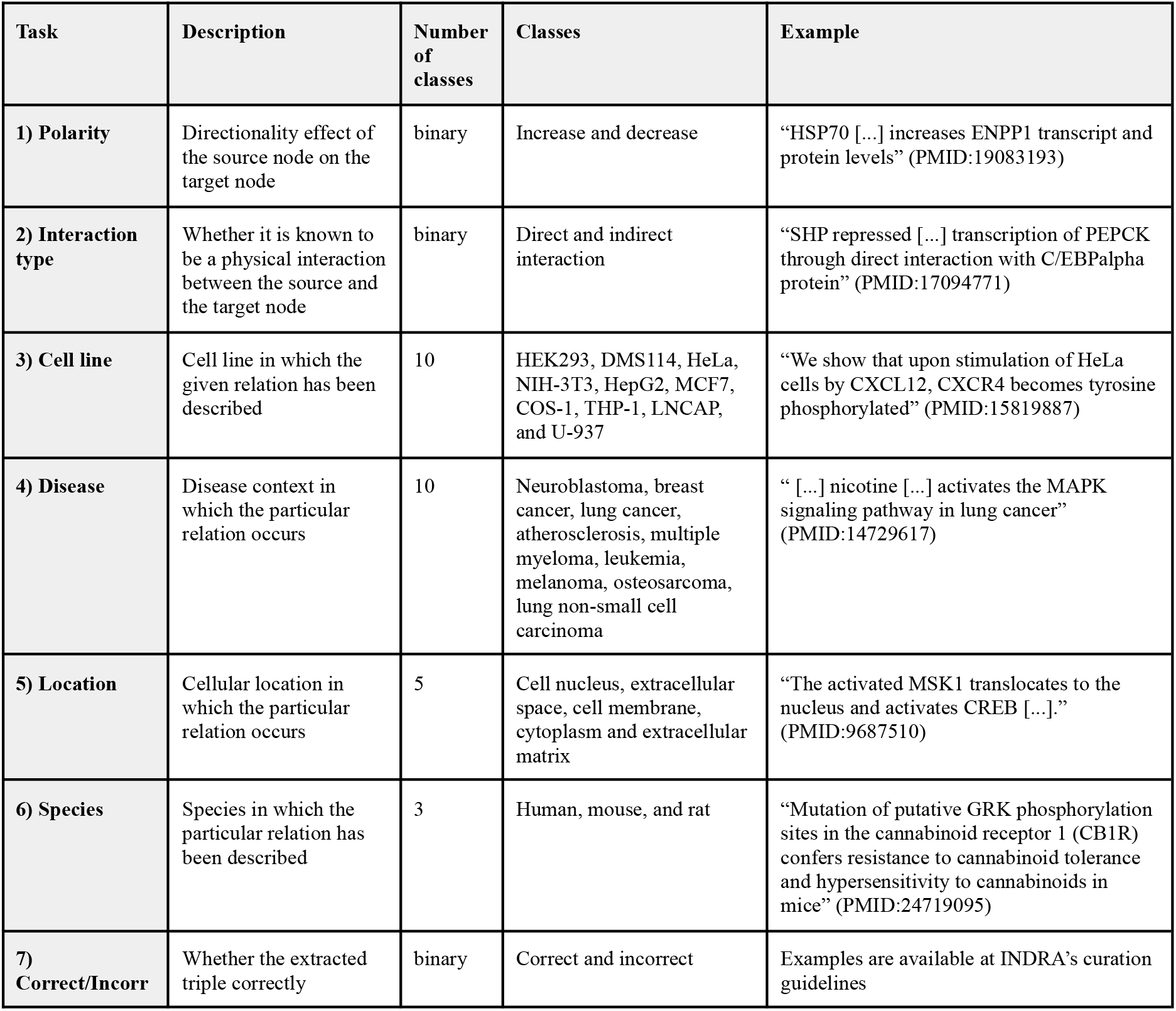
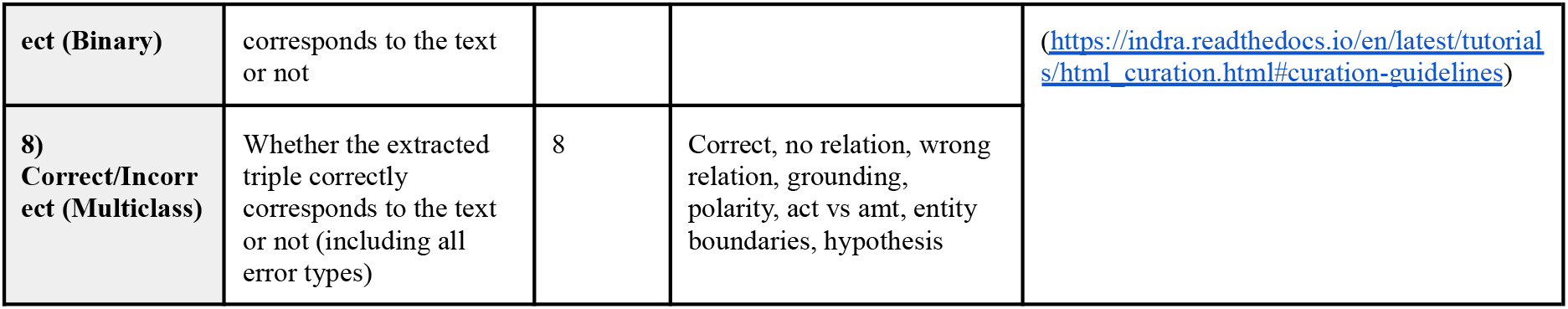
Overview on the fine-tuning classification tasks. While the two binary tasks (i.e., the polarity and interaction type tasks) intend to evaluate the models’ abilities to classify the relation type of the triple, the other four tasks deal with the classification of different types of contexts in which a given triple can appear in. Finally, the two tasks aim at predicting whether the triple has been correctly extracted from the text evidence.

The performance of the models was evaluated on all eight classification tasks via a five-fold cross-validation procedure using weighted F1-scores (i.e., averages of the class-specific F1-scores weighted by number of true instances per class). In order to train and evaluate all three models on the same cross-validation splits, we created the splits deterministically. All models were fine-tuned for five epochs on the training data using a batch size of 16 and the AdamW (Loshchilov and Hutter, 2019) optimizer with a linearly decreasing learning rate initially set to 5*10^−5^.

In addition to the proposed baselines, we introduced two ablated variants of the STonKGs model in order to analyze the effect of certain model design choices on the fine-tuning tasks:

1. **Less training steps:** We created two versions of the STonKGs model, STonKGs_150k_ and STonKGs_300k_, which were pre-trained for 150,000 and 300,000 steps (i.e., updates of the weights), respectively. More specifically, this was achieved through model checkpointing (i.e., STonKGs_150k_ is an interim checkpoint of STonKGs_300k_). In doing so, we were able to observe the effect of reducing the number of training steps on the model performance in the fine-tuning procedures.
2. **No NSP objective:** Since the effectiveness of the NSP task for pre-training has been questioned (see Liu *et al*., 2019), we decided to design a variant of STonKGs_150k_ (termed STonKGs_NO NSP_) that only uses the MLM and MEM training objectives. In result, this ablation measures whether the learned distinction between associated and randomly coupled text-triple pairs has an effect on fine-tuning task performances.

### 2.4. Implementation details

Both the NLP-baseline as well as STonKGs are implemented using the HuggingFace transformers library (v.4.6.1). More specifically, the NLP-baseline was initialized using the *dmis-lab/biobert-v1*.*1* BioBERT model available at the HuggingFace model hub. For STonKGs, we leveraged the *BertForPreTraining* class as a basis, and modified its prediction heads and forward pass function. STonKGs was pre-trained on 4x NVIDIA A100 40GB Tensor Core GPUs. The pre-training procedure took 284.18h (11.84 days) and 568.35h hours (23.68 days) for STonKGs_150k_ and STonKGs_300k_, respectively. Finally, to set up the KG-baseline, we employed the nodevectors library (v.0.1.23) for learning the random walk-based embedding sequences, and built a PyTorch Lightning (v.1.2.3) model on top. We trained our random walk-based embedding sequences on a symmetric multiprocessing (SMP) node with four Intel Xeon Platinum 8160 processors and 1.5TB RAM.

## 3. Results

### 3.1. Benchmarking

In order to analyze the differences in performance across the models in our benchmark setting, it is important to understand the information that is exploited by each baseline model. While the KG-baseline aims to represent topological node information, the NLP-baseline leverages the unstructured textual information underlying the relations between the extracted named entities (e.g., “*Rosiglitazone directly increases Pdk4 transcriptional-levels in mice*”). In our benchmark, six of the classification tasks consisted of predicting both type and context for each relation (e.g., a specific biological interaction is observed in a specific disease or species). Thus, the NLP-baseline seems more suited for these tasks compared to the KG-baseline, since the information could explicitly be stated in the evidence itself. Indeed, this is confirmed by our results, where we observed a better performance of the NLP-baseline over the KG-baseline across all tasks. Additionally, our proposed KG-baseline is limited by the use of static embeddings, as opposed to the transfer learning paradigm applied in both Transformer-based models (i.e., the NLP-baseline and STonKGs), which is based on fine-tuning the entire model architecture on given task-specific data. Below, we analyze the performances of the three presented models, as well as the ablated versions, across our proposed benchmark **(Table 2)**.

**Table 2.** Benchmark comparison of the baseline models and ablation variants of STonKGs on the chosen classification tasks. Performance is measured as the average F1-score across the 5 cross-validation splits. While the absolute performance gains are calculated based on the difference between the best STonKGs variant and the best baseline (i.e., the NLP baseline), the relative performance gains are obtained by dividing that difference by the f1-score of the best baseline and expressing the value as a percentage: 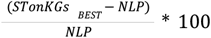.

| Model | Relation type classification task |  | Context annotation classification task |  |  |  | Correct/Incorrect classification task |  |
| --- | --- | --- | --- | --- | --- | --- | --- | --- |
|  | 1) Polarity | 2) Interaction type | 3) Cell line | 4) Disease | 5) Location | 6) Species | 7) Binary | 8) Multiclass |
| NLP-Baseline (BioBERT) | **0.940** | 0.991 | 0.238 | 0.214 | 0.397 | **0.865** | 0.911 | 0.881 |
| KG-Baseline (node2vec) | 0.448 | 0.945 | 0.020 | 0.030 | 0.295 | 0.670 | 0.708 | 0.446 |
| STonKGs<sub>300k</sub> | 0.930 | **0.995** | 0.252 | **0.248** | **0.405** | 0.860 | 0.977 | **0.964** |
| STonKGs<sub>150k</sub> | 0.931 | **0.995** | 0.256 | 0.240 | 0.404 | 0.860 | **0.978** | 0.963 |
| STonKGs<sub>NO NSP</sub> | 0.918 | 0.992 | **0.261** | 0.236 | 0.401 | 0.857 | 0.977 | 0.960 |
| Absolute performance gain | -0.009 | +0.004 | +0.023 | +0.034 | +0.008 | -0.005 | +0.067 | +0.083 |
| Relative performance gain | -0.96% | +0.40% | +8.81% | +15.89% | +2.02% | -0.58% | +7.35% | +9.42% |

Firstly, we focus on the four more challenging classification tasks (i.e., those containing more than five classes), namely, tasks 3-5 and 8 **(see Table 2**), where we could observe that STonKGs considerably outperformed both baselines. Here, STonKGs achieved between 0.01 and 0.08 larger F1-scores compared to the NLP-baseline. Compared to the KG-baseline, these differences were even larger resulting in F1-scores about 0.1-0.52 higher for STonKGs. Specifically for the cell line and disease tasks (task 3 and 4), the KG-baseline failed to predict the correct entity class among the ten possible classes, which was not the case for task 5 and 8, which both contain a lower number of classes. This suggests that it is particularly challenging for the KG-baseline to perform well across an increasing number of classes. On the other hand, when looking at the remaining four classification tasks containing with only two or three classes (i.e., task 1-2 and 6-7), we observe that both the NLP-baseline as well as STonKGs result in higher F1-scores than the KG-baseline. However, while STonKGs clearly outperforms the NLP-baseline with a difference of 0.067 on task 7, there are only minimal differences between the two models across tasks 1-2 and 6. While the NLP-baseline leads to a 0.009 and 0.005 improvement on the polarity and species tasks, STonKGs achieves a 0.004 F1-score improvement on the interaction task.

Interestingly, the KG-baseline is only slightly worse than the other two models on the interaction type and species classification tasks. On the polarity task, however, its performance is similar to a random classifier. The relative increase in performance of the KG-baseline on the interaction type task compared to the polarity task can be attributed to imbalanced associations between nodes and class labels (e.g., a given node might be exclusively present in indirect interactions) **(see Supplementary Figure 3)**. Moreover, in the case of species classification, the good performance of the KG-baseline is not surprising as the nodes in the INDRA KG can indirectly encode species information. For instance, protein nodes corresponding to the same ortholog gene are represented by species-specific identifiers (e.g., HGNC:PRKCG (human) and UP:P63319 (rat)).

When comparing STonKGs_300k_ and STonKGs_150k_, there is no significant difference in model performance. This is not surprising given the already low loss exhibited by STonKGs_150k_ and the minor further reduction of the loss in STonKGs_300k_ **(Supplementary Figure 4)**. Moreover, the STonKGs_NO NSP_ model resulted in slightly lower performances than the STonKGs_150k_ model on almost all the evaluation tasks (apart from task 3), thus, suggesting that the NSP training objective is potentially beneficial for the overall pre-training procedure.

### 3.2. STonKGs and Applications

The source code and trained models are respectively available at https://github.com/stonkgs/stonkgs and https://github.com/stonkgs/results. The documentation is available at https://stonkgs.readthedocs.io/. The pre-trained STonKGs model can be downloaded from the HuggingFace model hub (https://huggingface.co/stonkgs/stonkgs-150k).

To demonstrate the generality of the pre-trained STonKGs model, we fine-tuned it on INDRA-independent text-triple pairs specific to two neurodegenerative indication areas (i.e., Alzheimer’s disease, Parkinson’s disease) (Domingo-Fernández *et al*., 2017) (presented in the **Supplementary Text 2**). Furthermore, the fine-tuned STonKGs models, which are also released, can also be used to automatically annotate text-triple pairs with respect to the defined classes for each fine-tuning task (e.g., human, mouse, and rat for the species context annotation task); thus, facilitating automatic annotations of biomedical KGs in a variety of contexts.

## 4. Discussion

In this work, we introduced STonKGs, a multimodal Transformer trained on millions of text-triple pairs from biomedical literature assembled by INDRA. We demonstrate the utility of our approach in a benchmark consisting of eight fine-tuning tasks. Here, STonKGs outperformed two baseline models, which were trained solely on either text or KG data, on the majority of the benchmark tasks. Each of the eight fine-tuning tasks represents a different classification problem with a specific biological use-case, hence confirming the generalizability of our proposed transfer learning approach. In addition to the benchmark, we conducted further ablation studies to measure the influence of the number of training steps and the NSP training objective on the overall performance of STonKGs. Finally, the source code and the pre-trained model are available at https://github.com/stonkgs, enabling to leverage both the pre-trained STonKGs model as well as the overall model architecture for a variety of additional ML-based tasks that use text and KG data.

There exist some limitations to our proposed STonKGs model. Firstly, while we have trained STonKGs on a novel and comprehensive KG that has not been utilized by any other Transformer-based model before, the INDRA KG is comparatively smaller than other large-scale available non-biomedical KGs such as Wikidata (Vrandečić and Krötzsch, 2014) and DBpedia (Bizer *et al*., 2007). This is mainly caused by the challenging tasks of recognizing biological entities and extracting their relations, given the ambiguity and complexity of biomedical jargon. Furthermore, INDRA aims at high precision and the employed extraction process focuses on high quality rather than completeness. This impacted the text-triple pairs present in the fine-tuning datasets (i.e., some of them contain only several thousand text-triple pairs). Secondly, one characteristic property of the INDRA KG is that its textual evidences have been extracted on sentence level, consequently they are shorter in length compared to text sequences used in other Transformer-based LMs (e.g., Devlin *et al*. (2019) and Zaheer *et al*. (2020)). Given the complexity of biological scientific literature, the contextual representations learned by STonKGs could benefit from longer sequences (i.e., the context of a given triple is often mentioned in the surrounding sentences). Thirdly, while we have generated the node embeddings based on node2vec, other more sophisticated models such as Graph Convolutional Networks (GCNs) and Graph Attention Networks (GATs) (Ji *et al*., 2021) could be employed. However, this is practically infeasible due to the computational complexity required given the size of our KG. Furthermore, there are two advantages of using node2vec as opposed to employing other models: i) node2vec scales well for large-scale KGs, and ii) sequential input is implicitly generated by using random walks for each node. Another limitation is the absence of an optimization procedure for hyperparameters such as the batch size or the learning rate of STonKGs due to the run time implications (i.e., pre-training required several weeks, and running all benchmark tasks for STonKGs took more than a day). However, we demonstrated the effectiveness of STonKGs using the standard hyperparameters from the original BERT model. Finally, there are at least reasons why we could not include other KG-extended Transformers (i.e., Zhang *et al*. (2019), He *et al*. (2020), and Fei *et al*. (2020)) in our benchmark setting: i) these models require entity linking between text and KG nodes **(see Introduction)**, and ii) our benchmark is specifically designed to evaluate the performance of the models in classifying context and relation type information, which is not covered in benchmarks of other approaches.

Although we have demonstrated a proof-of-concept of our methodology across a variety of classification tasks, we would like to mention possible future improvements of STonKGs. Firstly, the STonKGs pre-training procedure could potentially benefit from an even larger corpus of text-triple pairs. Due to our proposed transfer learning setting, additional corpora of text-triple pairs can be flexibly fed into the model by continuing the pre-training procedure. Secondly, while we have proposed a novel method to generate contextualized graph embedding sequences based on random walks from node2vec, more powerful KGE models could be potentially adapted to generate sequential input embeddings as well. Thirdly, to maximize information gain, textual descriptions of the KG nodes could be added to the model in a straightforward manner. Finally, an in-depth analysis of the attention weights between the text tokens and KG nodes used in STonKGs could reveal valuable insights about the interdependencies between the two modalities.

## Authors’ Contributions

HB conceived and designed the study together with DDF. HB implemented the methodology and conducted the experiments supervised by DDF. CTH, CB, BG, and JB assisted in defining the experiments. CTH, BG, and JB provided and processed the data. MHA, ATK, BG, JB, and DDF acquired the funding. HB, CB, and DDF wrote the manuscript. CTH, BG, JB, and PGP reviewed the manuscript.

