## Supplementary File for "STonKGs: A Sophisticated Transformer Trained on Biomedical Text and Knowledge Graphs"

#### Supplementary Text

##### 1. INDRA Knowledge Graph

The INDRA KG was generated from a collection of corpus described in the Methods section. The KG contains several relations connecting nodes representing biological entities from a variety of modalities. These relations (edges/triples) have been extracted from INDRA (http://www.indra.bio/) (Gyori *et al.*, 2017) from the textual evidence found in the corpus using an ensemble of natural language processing systems. INDRA also contains triples from structured databases such as SIGNOR (https://signor.uniroma2.it/). While some structured sources provide evidence text corresponding to relations, others (e.g., BioGRID) do not.

The INDRA KG was generated on 28.04.2021 and comprised 35,150,093 text-triple pairs (also known as statements). To guarantee the quality of each triple in the KG, we filtered out any triple that i) contained any ungrounded node (i.e., nodes that could not be normalized to an standardized ontology), or ii) did not contain textual evidence, or iii) that was not part of the largest component of the graph, resulting in 13,609,994 filtered text-triple pairs. These filtering steps were conducted after loading the INDRA statements using PyBEL (Hoyt *et al.*, 2018). Additionally, this filtering step ensures that the vocabulary size of the KG stays in the same order of magnitude as the token vocabulary size of the BERT model. This filtered version of the INDRA KG is summarized in **Supplementary Figure 1.**

In this filtered version, there are four relation types: increases (INC), decreases (DEC), directly decreases (DIR-DEC), and directly increases (DIR-INC). Following, we generated the required splits for pre-training and each of the eight fine-tuning tasks detailed in the manuscript. To avoid data leakage, we generated mutually exclusive sets of triples for pre-training and fine-tuning. Note, however, that the fine-tuning datasets themselves are not mutually exclusive, since a given triple might contain multiple annotations (e.g., both a species and a cell line annotation), which might be used for multiple tasks. For each annotation type task **(task 3-6)**, its respective data split consisted of all available text-triple pairs containing that particular annotation, as summarized in **Supplementary Table 1**. On the other hand, for both the interaction type and polarity tasks **(task 1 and 2)**, we used a shared subset of 78,979 text-triple triples of protein-protein interactions (i.e., complexes, biological processes, and reaction nodes are excluded) as a basis for the respective splits, consisting of between 18,638 and 20,811 text-triple pairs per relation type. The rationale behind the choice of the subset size is two fold. First, we aimed at maximizing the number of text-triple pairs used for the pre-training procedure. Second, it allowed us to train on a roughly evenly distributed number of classes in both classification tasks and to avoid class imbalance. This is possible since any of the four relations present in the KG can be used in both tasks **(see Supplementary Table 3).** Given the relatively low number of text-triple triples with the DIR-DEC relation type (n=18,638) compared to the rest , the chosen split size allows us to maintain an equal number of classes in the fine-tuning task while also using a sufficient number of text-triple triples from this class in the pre-training procedure. Lastly, the two correct/incorrect classification tasks **(task 7 and 8)** consist of 12,836 and 12,611 text-triple pairs that have been manually curated for machine learning applications (https://indra.readthedocs.io/en/latest/tutorials/html_curation.html#curation-guidelines). This curated dataset consists of assembled INDRA statements/triples and a label assigned by a curator indicating whether the statement/triple has been correctly extracted from the evidence or not. In case, there was an error in the extraction, the type of error is also labelled, which enabled us to conduct two different classification tasks: i) correct vs. incorrect, and ii) correct and all incorrect types as independent labels.

Finally, we would like to mention that for all fine-tuning task datasets, we removed duplicate text-triple pairs (i.e., sets of text-triple pairs that contain the same text evidence) to prevent that a specific evidence is present in both training and test splits. Additionally, we also excluded text-triple pairs containing triples with nodes that are not present in the pre-trained node2vec model, since they cannot be mapped to an embedding representation.

##### 2. Custom Application of STonKGs Classifying Text-triple Pairs Based on Neurodegenerative Disorders

The starting point for the neurodegenerative application case is an independent (i.e., coming from a different data source than INDRA) fine-tuning dataset (Domingo-Fernández *et al.*, 2017), consisting of 2,264 annotated text-triple pairs from the Alzheimer's and Parkinson's, and epilepsy BEL graphs at https://github.com/neurommsig/neurommsig-knowledge. The initial annotations contained three classes representing context annotations for different neurodegenerative diseases: Alzheimer's Disease (AD), Parkinson's Disease (PD) and epilepsy. Out of these text-triple pairs, 695 many had source and target nodes that were both contained in the INDRA KG, hence, this subset was used for the remainder. The epilepsy class was contained in less than 1% of the labels in this subset, therefore it was left out, resulting in a binary classification task (AD vs PD). Based on initializing the pre-trained $\text{STonKGs}_{300k}$ model with a sequence classification head and using the same five-fold cross-validation procedure and metrics as for the other classification tasks **(see Evaluation)**, we achieved a mean F1-score of 0.898.

### Supplementary Figures

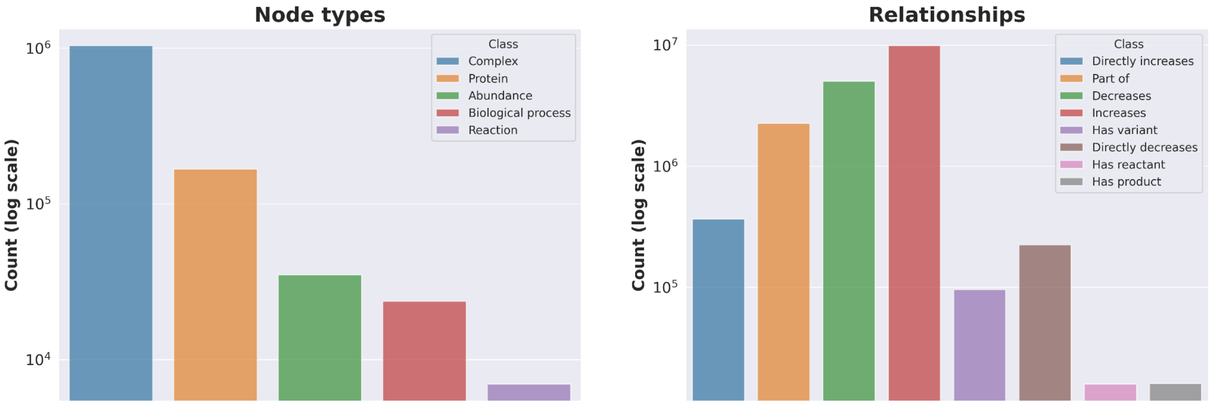


**Supplementary Figure 1.** Class distributions of node and relationship types in the filtered dataset (prior to splitting it into pre-training and fine-tuning partitions).

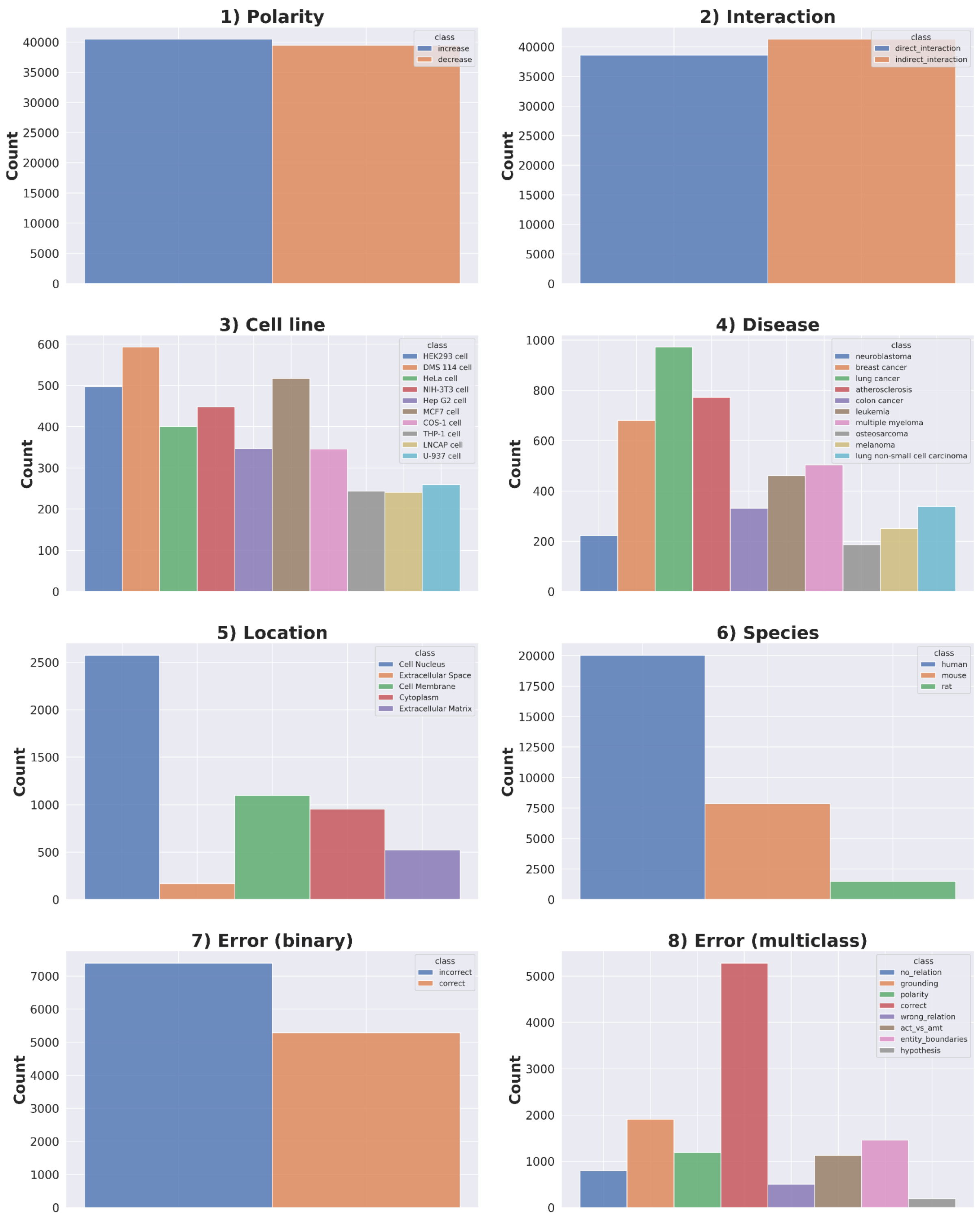


**Supplementary Figure 2.** Class distributions of all fine-tuning tasks, based on the task-specific datasets, which have been previously filtered for their respective majority classes.

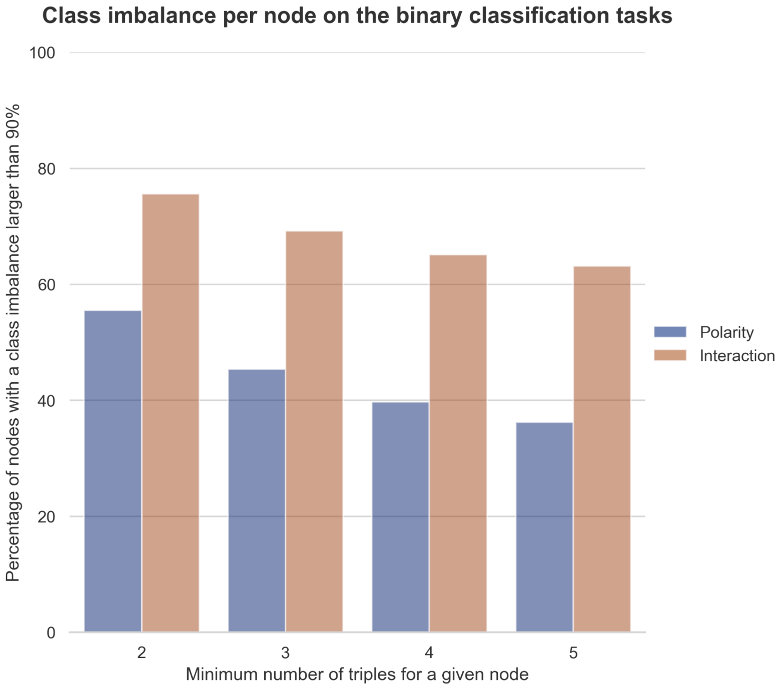


**Supplementary Figure 3. Class imbalance for nodes present in multiple triples in the polarity and interaction tasks.** The bar charts show the proportion of nodes that are imbalanced to a specific class in more than 90% of triples that the node appears in. This proportion is calculated based on a minimum threshold that specifies the minimum number of triples in which a given node is present (x-axis). We can observe that the majority of the nodes in the interaction type classification task exhibit such a bias, compared to a much smaller proportion in the polarity task. In fact, 19 out of the 25 nodes with the most triples (>800 triples) in the task-specific dataset exhibit the previously defined class-specific bias. Thus, it can be expected that in the training procedure of the KG-baseline, the model learns patterns for the given set of nodes.

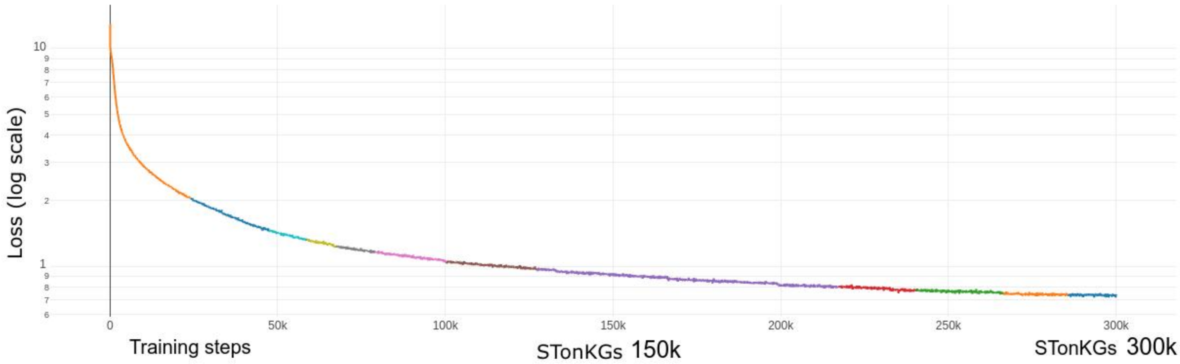


**Supplementary Figure 4. Loss curve for the pre-training process of $STonKGs_{150k}$ and $STonKGs_{300k}$, consisting of the sum of the losses of the MLM, MEM and NSP objectives.** The x-axis indicates the number of training steps, and the y-axis displays the loss at a log scale. Moreover, the different colors of the loss curve represent different checkpoints with which both models were trained until 150,000 or 300,000 training steps were reached for $STonKGs_{150k}$ and $STonKGs_{300k}$, respectively.

### Supplementary Tables

| Reading System | Reference |
|---|---|
| **REACH** | Valenzuela-Escárcega *et al.* 2018 |
| **TRIPS** | Allen *et al.* 2015 |
| **Sparser** | https://github.com/ddmcdonald/sparser |
| **MedScan** | Novichkova *et al.* 2003 |
| **TEES** | Björg, 2014 |
| **ISI** | Garg *et al*. 2016 |
| **Geneways** | Rzhetsky *et al*. 2004 |
| **RLIMS-P** | Torii *et al.* 2014 |
| **Eidos** | https://github.com/clulab/eidos |
| **GNBR** | https://zenodo.org/record/3459420 |

**Supplementary Table 1**. **List of reading systems used by INDRA.**

| Classification task | Number of triples fine-tuning split | Number of classes | Ontology |
|---|---|---|---|
| **1) Cell line** | 3,760 | 10 | Cell Line Ontology |
| **2) Disease** | 4,586 | 10 | Disease Ontology |
| **3) Location** | 5,223 | 5 | MeSH |
| **4) Species** | 21,765 | 3 | NCBI Taxonomy |
| **5) Interaction type** | 78,979 | 2 | - |
| **6) Polarity** | 78,979 | 2 | - |
| **7) Correct/Incorrect (Binary)** | 12,836 | 2 | - |
| **8) Correct/Incorrect (Multiclasss)** | 12,611 | 8 | - |

**Supplementary Table 2**. **Number of triples and classes used for each fine-tuning task.** The distribution of each of the classes in each fine tuning task are displayed in **Supplementary Figure 1**.

| Hyperparameter | NLP baseline (BioBERT) | KG baseline (node2vec) | STonKGs |
|---|---|---|---|
| **Initial learning rate** | $10^{-4}$ | 0.025 (for the internal word2vec algorithm) | $10^{-4}$ |
| **Learning rate scheduler** | linear (i.e., linearly decreasing), warmup over the first 10,000 steps | linear (i.e., linearly decreasing), no warmup | linear (i.e., linearly decreasing), no warmup |
| **Optimizer** | Adam | Stochastic Gradient Descent (for the internal word2vec algorithm) | AdamW |
| **Batch size (before gradient accumulation)** | 192 | 10,000 (for the internal word2vec algorithm) | 64 |

| | | | |
|---|---|---|---|
| **Gradient accumulation steps** | - | - | 8 |
| **Effective batch size** | 192 | 10,000 (for the internal word2vec algorithm) | 512 |
| **Maximum sequence length** | 512 | - | 512 |
| **Training steps** | 1,000,000 | - | 300,000 ($STonKGs_{300k}$) or 150,000 ($STonKGs_{150k}$) |
| **Epochs** | - | 1 (for node2vec), 4 (for the internal word2vec algorithm) | - |
| **Half-precision floating-point format** | No | No | Yes |
| **Embedding dimension** | 768 | 768 | 768 |
| **Random walk length** | - | 127 | - |
| **Window size** | - | 3 | - |
| **Number of negative samples** | - | 5 | - |
| **Return weight** | - | 1.0 | - |
| **Neighbor weight** | - | 1.0 | - |

**Supplementary Table 3. Summary of the central hyperparameters used in pre-training for each of the three models employed in this approach. The hyperparameters for BioBERT are directly taken from Lee *et al.* (2020).**

| **Relation** | **Interaction type** | **Polarity** |
|---|---|---|
| **Increases (INC)** | Indirect | Up |
| **Decreases (DEC)** | Indirect | Down |
| **Directly Increases (DIR-INC)** | Direct | Up |
| **Directly Decreases (DIR-DEC)** | Direct | Down |

**Supplementary Table 4. Different use of the four different relation types for the interaction type and polarity fine-tuning tasks.** In the interaction type task, INC and DEC are labelled as “indirect”, whereas DIR-INC and DIR-DEC are labelled as “direct”. For the polarity task, INC and DIR-INC are grouped together for up regulation, whereas DIR and DIR-DEC are associated with the label for down regulation.
